# A pilot study of behavioral, environmental, and occupational risk factors for chronic kidney disease of unknown etiology in Sri Lanka

**DOI:** 10.1101/837393

**Authors:** Jake M. Pry, Wendi Jackson, Ruwini Rupasinghe, Guneratne Lishanthe, Zied Badurdeen, Tilak Abeysekara, Rohana Chandrajith, Woutrina Smith, Saumya Wickramasinghe

## Abstract

Chronic kidney disease of unknown etiology (CKDu) was first recognized in Sri Lanka in the early 1990s, and since then it has reached epidemic levels in the North Central Province of the country. The prevalence of CKDu is reportedly highest among communities that engage in chena and paddy farming, which is most often practiced in the dry zone including the North Central and East Central Provinces of Sri Lanka. Previous studies have suggested varied hypotheses for the etiology of CKDu; however, there is not yet a consensus on the primary risk factors, possibly due to disparate study designs, sample populations, and methodologies. The goal of this pilot case-control study was to evaluate the relationships between key demographic, cultural, and occupational variables as risk factors for CKDu, with a primary interest in pesticide exposure both occupationally and through its potential use as an ingredient in brewed kasippu alcohol. A total of 56 CKDu cases and 54 control individuals were surveyed using a proctored, self-reported questionnaire. Occupational pesticide exposure and alcohol consumption were not found to be significant risk factors for CKDu. However, a statistically significant association with CKDu was observed with chewing betel (OR: 6.11, 95% CI: 1.93, 19.35), age (OR: 1.07, 95% CI: 1.02, 1.13), owning a pet dog (OR: 3.74, 95% CI: 1.38, 10.11), water treatment (OR: 3.68, 95% CI: 1.09, 12.43) and pests in the house (OR: 5.81, 95% CI: 1.56, 21.60). The findings of this study suggest future research should focus on practices associated with chewing betel, potential animal interactions including pests in the home and pets, and risk factors associated with water.

**AUTHOR SUMMARY:** Since a new variant of chronic kidney disease was acknowledged in the early 1990s among those in the agricultural community of Sri Lanka, especially rice farmers, the research community has searched for causes of what has become known as chronic kidney disease of unknown etiology or CKDu. Previous studies have focused on heavy metals in the environment as they are known to be toxic to the kidneys however, a proverbial “smoking gun” has yet to be identified. Understanding that the causes is potential multifactorial we implemented a pilot case-control study using a One Health methodology administering a comprehensive interview to assess environmental, animal, and, human exposures that may be contributing to the diagnosis of CKDu. We found statistically significant odds ratio among those that reported having a pet dog, chewing betel (a traditional preparation or various ingredients wrapped in a betel leaf inserted between the teeth and cheek), pests in the home, treating drinking water, and older age. These results serve to guide further hypothesis generation regarding mechanisms behind associated exposures from infectious diseases such as hantavirus and leptospirosis to food preparation through boiling drinking water in aluminum vessels and oral pesticide exposure linked to betel preparation.

## INTRODUCTION

There has been a notable increase in the recognized incidence of chronic kidney disease (CKD) around the world [1]. Kidney disease has moved from 27^th^ most common cause of death in 1990, to 18^th^ in 2010 and has come to be considered a global public health problem causing high morbidity, mortality, and financial burden [2-4]. Global prevalence of CKD is estimated to range between 8% and 16%, and differs substantially across developed and developing countries [3, 5]. Although diabetes mellitus and hypertension remain the leading causes of CKD, in recent years a different form of CKD has reached epidemic levels, devastating rural communities in the dry zone of Sri Lanka [6, 7]. The recognition of endemic CKD in the dry zone in the 1990s coincided with the development of the rural healthcare system, which improved access to clinics by affected individuals. Since that time, the dry zone has seen a disproportionate increase in cases of CKD compared to the rest of the country [8]. Existing studies describe the majority of these CKD patients as not having hypertension or diabetes mellitus, two of the major risk factors for CKD. It has therefore, been defined as a distinct condition: CKD of unknown or uncertain etiology (CKDu). Similar chronic kidney disease hotspots have been recognized among farmers in Central America (Nicaragua and El Salvador) and South Asia [9, 10].

Approximately 2.5 million people live in the subset of Sri Lankan provinces where CKDu is most common [11, 12]. Cases of this disease predominate in the Medawachchiya, Wilgamuwa, Nikawewa, and Girandurukotte regions of the dry zone. Studies have shown the highest prevalence of CKDu among 30-60 year old men engaged in chena or rice farming, and estimate a total of 20,000 (approximately 0.8% population) affected in the North Central Province [8, 13].

The epidemic of CKDu in the dry zone is burdening the rural healthcare system and impacting agricultural productivity due to a reduction in the available labor force when CKDu patients are too ill to work [14-16]. Due to the irreversible and progressive nature of CKD, most patients require long-term dialysis since renal transplants are not commonly available. For these reasons, there is a need to determine the risk factors associated with CKDu to control and attenuate the incidence of new CKDu cases. A growing body of evidence suggests that CKDu is multi-factorial, making it difficult to identify individual risk factors and potential interactions involved in pathogenesis [7, 12, 17-20]. Recently, various heavy metal agents such as cadmium, arsenic agrochemicals, aluminum, and fluoride, as well as infectious diseases such as leptospirosis have been considered for association with CKDu [21-26].

Collaboration between researchers at the University of Peradeniya in Sri Lanka, the University of California, Davis (UCD) in the United States, and Sri Lankan stakeholders in CKDu-endemic areas were involved in this pilot study. The driving hypothesis for this study is that alcohol consumption and/or pesticide exposure are associated with CKDu as a health outcome. In addition, it is recognized that relationships between key demographic, cultural, and occupational variables may play a role in CKDu health outcomes [27-31].

## MATERIALS AND METHODS

A pilot case-control study was conducted in Sri Lanka from July-October 2015. The study population was comprised of individuals (cases and controls) who resided in the North Central Province (NCP) or Uva Province (UP) and sought medical care at Girandurukotte district hospital (UP) or Medawachchiya clinic (NCP). The population in both the NCP and UP is approximately 1.2 million, with women making up the slight majority (51%) [13]. The majority of people in both provinces are Sinhalese-speaking and resides in the rural sector where they engage in farming (chena, rice). The NCP has the highest recorded prevalence of CKDu cases in Sri Lanka and is located in the country’s dry zone. Uva Province is in the intermediate zone adjacent to the dry zone, with a lower prevalence of CKDu cases compared to the NCP.

In order to test the hypothesis that there is a relationship between alcohol consumption and CKDu diagnosis, and pesticide exposure and CKDu, in endemic areas of Sri Lanka, a questionnaire survey was developed [See Additional File 1]. The survey tool encompassed a wide range of exposures to capture potential unknown confounders, including exposures suggested by local CKDu working groups at the University of Peradeniya. Individuals meeting the CKDu case definitions as well as a comparison (control) population from the same endemic region were invited for participation in the survey.

### Case Definition

Individuals diagnosed with definite or probable CKDu by a nephrologist at the Girandurukotte regional hospital (GRH) or Medawachchiya renal clinic (MRC) made up the population from which study cases were selected. An individual was considered a definite CKDu case if creatinine levels were elevated and in subsequent renal biopsy the finding was predominant tubular interstitial nephritis. A probable CKDu case was defined as persistent renal dysfunction for more than 3 months after excluding known causes including hypertension, diabetes mellitus, any other known renal diseases. This methodology is consistent with clinic/hospital programs and represents the CKDu process of diagnosis in the region. Controls were chosen based on negative results for CKDu from population screening records at GRH or MRC.

### Recruitment

All controls were recruited using CKDu screening results within the past three years. These CKDu negative were invited via post (hard copy letter) to return to the healthcare facility associated with the previous screening to take part in a survey. Participation in the survey was optional. All cases were recruited from Girandurukotte regional hospital or Medawachchiya renal clinic.

### Sample Size Calculation

The total sample size calculated for this pilot case-control study was 110, comprising 1:1 cases to controls. The target sample size of 110 individuals was calculated based on a power of 80% (β= 0.2), 95% confidence (α= 0.05), and a minimum effect size of 3.0. This relatively large effect size was considered in the exploratory study in order to identify preliminary exposures strongly associated with the outcome. An estimate of 26% was used for any reported alcohol consumption among controls for the sample size calculation [32, 33].

### Survey Design

Survey questions were designed by the research team in consultation with the resource personnel at the Centre for Research, Education, and Training on Kidney Diseases (CERTkID) in Sri Lanka prior to IRB approval, survey training, and interviews. The survey consisted of 138 questions structured as binary, categorical, ordinal, and open-ended across six categories: 1) agricultural information; 2) animal exposure; 3) water and nutrition; 4) alcohol consumption; 5) respondent demographics; and 6) family and past medical history.

The agricultural information section included questions related to farming practices and agrochemical usage. Information on ownership and health of livestock and pets, presence of pest animals and wild life were collected in section two of the survey. In the water and nutrition information section, sources for drinking, cooking, and bathing water were assessed, along with participant practices regarding water treatment prior to use. The alcohol consumption information section contained questions on type of alcohol consumed, betel chewing and smoking status. Alcohol consumption was assessed in two ways: a binary question was asked first on whether the participant had ever consumed alcohol (if yes, what type and frequency) and second, whether the participant believed that alcohol was a problem in their village. The respondent demographics section contained questions pertaining to level of education and family income. To assess a potential genetic component of CKDu, participants were asked whether their spouses were close blood relatives and family history of CKDu, hypertension, and diabetes mellitus was also recorded. Detailed survey and explanation of survey components are given in Supplementary information.

Survey data were collected by 12 trained graduate students associated with the University of Peradeniya. In addition, investigators were present at each site during data collection, allowing surveyors the opportunity for clarification as needed in real-time as interviews were conducted. Cases and controls participated voluntarily, and surveys were administered verbally in the mother tongue of the participants (Sinhala) after obtaining consent (see Supplementary Information for the English version of the consent form and survey questions). Survey question responses were recorded on paper copies of the questionnaire by the surveyor. Each interview took approximately one hour to complete.

The research protocol was designed according to the guidelines of the International Compilation of Human Research Standards (2015 edition) and approved by the University of California, Davis Institutional Review Board (#762486-2). Written consent of all study participants was obtained by signature or thumbprint after survey enumerators verbally read the consent statement in the appropriate language. The consent form was translated in both Sinhala and Tamil. The majority of survey interviews took place at medical clinics specializing in renal disease.

### Statistical Analyses

Logistic regression was used to evaluate risk factors for CKDu case status. Bivariate analysis was used to identify covariate with p-value ≤ 0.20 which was used to restrict consideration for the final model. Pearson’s correlation coefficient was used to identify covariate correlation at −0.5 ≤ *ρ* ≥ 0.5. Data analysis was completed using Stata IC 14 (StataCorp. 2015. Stata Statistical Software: Release 14. College Station, TX: StataCorp LP). Age was evaluated as a continuous predictor and others were assessed as binary or categorical. Multiple logistic regression analyses were performed using backwards stepwise selection to model the risk factors associated with the CKDu disease outcome of interest. Adjusted analysis was done to control for possible confounding by measured covariates. Statistical significance was assessed at the *α*= 0.05. Any observations missing data were restricted from the analysis dataset. All geospatial illustrations were prepared using QGIS v2.18 (Free Software Foundation Inc., Boston MA, USA) with administrative boundaries provided by GADM v2.8 (Center for Spatial Statistic, University of California Davis, USA, accessed November 2015).

## RESULTS

A total of 110 participants were included in the analysis; 56 met the case definition and 54 satisfied control criteria (Table S1). All participants resided in the CKDu endemic regions in Girandurukotte and Medawachchiya districts in Sri Lanka at the time of diagnosis. No missing data was identified in the analysis dataset. Participants had a mean age of 52.6 years (range = 25-80); there was a slight majority of males (60%) to females (40%). Most participants reported to be married, with about half reporting being married to a spouse that was a close blood relative and slightly over half reporting a family member having been diagnosed with CKDu (Table S1). Of the 110 study participants, half reported consuming any type of alcohol and the majority reported using some type of pesticide in their daily lives (insecticide, herbicide, in-home pesticide and/or fungicide) (Table S1).

The majority of participants (74) were residing in the Uva Province. Twenty-two participants resided in the North Central Province, five in the Eastern Province, two in the Central Province and one resided in the Northwestern Province. There were 8 (7.3%) individuals for whom reliable current residence information was not available due to survey legibility and standardization complications. Data regarding province in which the participant was born were collected (Fig 1). No participants reported birth outside of Sri Lanka.

**Figure 1.**
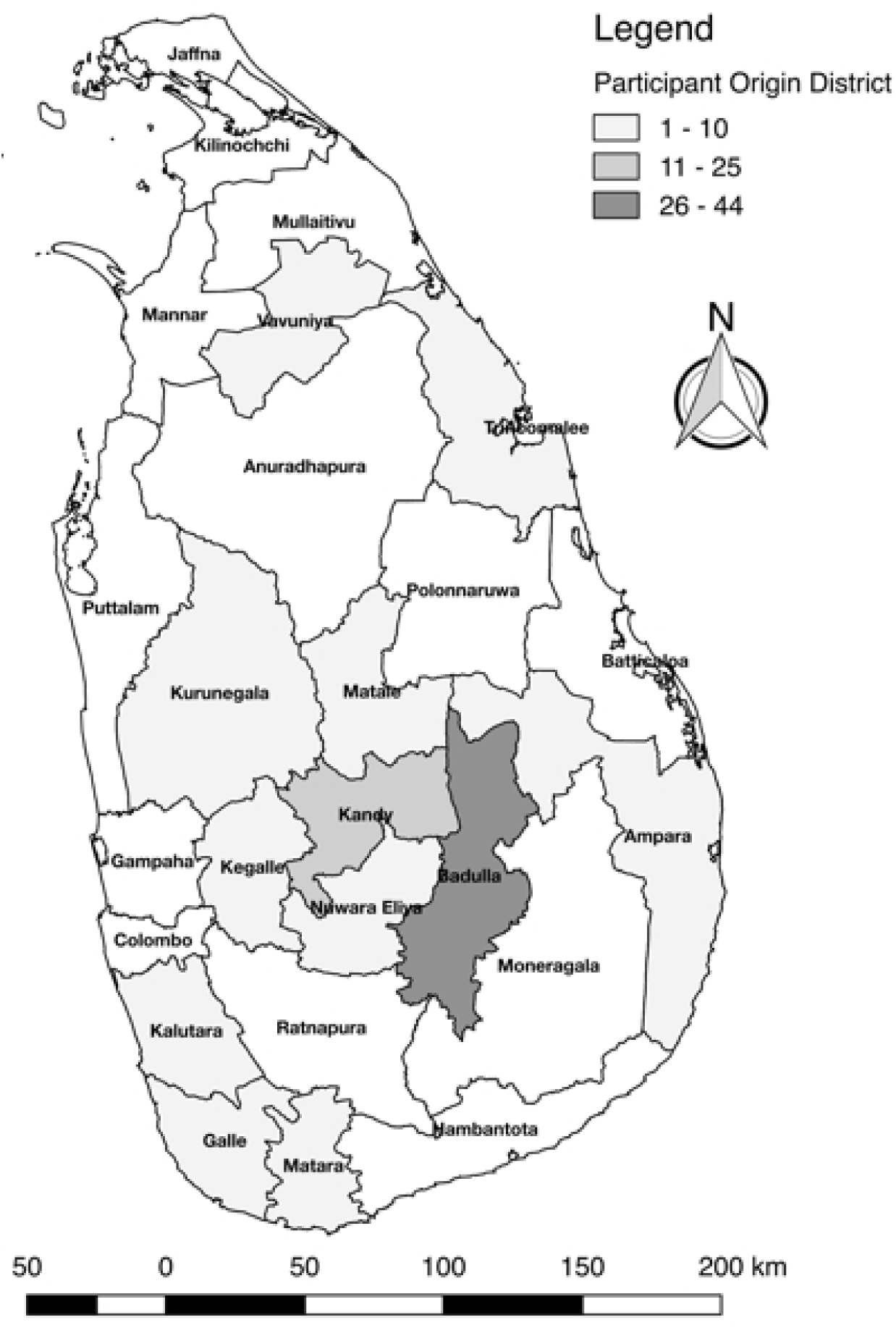
Map of Sri Lanka by district indicating participant’s birth district

After bivariate and correlational analyses, smoking unfiltered cigarettes and smoking cannabis were found to be highly correlated (correlation coefficient = 0.5141 > 0.5 threshold). Smoking unfiltered cigarettes was dropped from model consideration given a bivariate p-value higher than that of smoking cannabis (p-value = 0.186 and 0.112, respectively).

Sources of drinking water were surveyed and ‘dug well’ was the most common source (90.9%) of drinking/cooking water, with ‘rainwater collection containers’ being the second most common source. Drinking water was reported to be treated with routine methods such as boiling or filtering, or traditional methods such as placing igini (*strychnos potatorum*) seeds in the water source/well [34]. A subset of key population characteristics is reported in Table 1.

**Table 1:** Study Population Characteristics by Case-Control Status.

| *Factor* | *Level* | *Control* | *Case* | *p-value* |
| --- | --- | --- | --- | --- |
| *N* |  | 54 | 56 |  |
| Age, mean (SD)* |  | 49.5 (11.7) | 57.5 (9.6) | <0.001 |
| Gender* | Female | 27 (51) | 16 (29) | 0.017 |
|  | Male | 26 (49) | 40 (71) |  |
| Farming as Occupation* | No | 14 (28) | 3 (6) | 0.002 |
|  | Yes | 36 (72) | 51 (94) |  |
| Drinking Water Source | Dug Well | 48 (89) | 52 (93) | 0.47 |
|  | Rain Water | 6 (11) | 5 (9) | 0.7 |
| Treat Drinking Water | Drinking | 34 (63) | 41 (75) | 0.19 |
| Keep Livestock | Livestock | 17 (31) | 19 (34) | 0.78 |
| Smoking Status | Tobacco | 8 (15) | 14 (25) | 0.18 |
|  | Cannabis | 2 (4) | 7 (12) | 0.092 |
| Chew Betel* | Betel | 22 (41) | 40 (71) | 0.001 |
| Alcohol a Problem in Village | Not a Problem | 13 (27) | 11 (21) | 0.85 |
|  | Minor Problem | 17 (35) | 20 (38) |  |
|  | Moderate Problem | 10 (20) | 9 (17) |  |
|  | Major Problem | 9 (18) | 12 (23) |  |
| Alcohol Consumption | Any Alcohol | 22 (41) | 33 (59) | 0.056 |
|  | Arrack | 19 (35) | 27 (48) | 0.17 |
|  | Beer | 14 (26) | 16 (29) | 0.76 |
|  | Kasippu | 13 (24) | 18 (32) | 0.35 |
| Pesticide Use* | Any | 42 (78) | 52 (93) | 0.025 |
|  | Fungicide | 21 (44) | 29 (55) | 0.27 |
|  | Herbicide | 41 (76) | 51 (91) | 0.032 |
|  | Insecticide | 42 (78) | 44 (79) | 0.92 |

\*Significance attributed as p-value < 0.05

There is a significant difference between age, gender, occupation, chewing betel, pesticide use at the 95% confidence level. The cases were older, on average, by eight years than the control group with a gender imbalance finding males more often among the cases compared to controls. Cases cited farming as an occupation more often than controls (22% higher among cases), reported chewing betel (30% higher among cases), and used pesticides more often, specifically herbicide, (15% higher among cases).

Two multivariable models were constructed, an exposure of interest model and an exploratory model. The exposure of interest model forced inclusion of variables concerning alcohol consumption and pesticide exposure, as neither exposure of interest was found to be significantly associated with CKDu in bivariate analysis (Table 2). The final exposure of interest model included four variables and excluded one variable compared to the stepwise method used for the exploratory model (Table 3). The exploratory model (Table 3) included only risk factors significantly associated with CKDu status (P<0.05). The primary exposures of interest (alcohol consumption and pesticide exposure) were not found to be significant using a backward stepwise selection process. However, age – considered as a continuous variable (OR: 1.08, 95% CI 1.02, 1.13), chewing betel (OR 4.01, 95% CI 1.49, 10.81), keeping a pet dog (OR: 4.21, 95% CI 1.55, 11.48), and reporting pests in the home (OR: 3.96, 95% CI 1.21, 12.93) were significantly associated with CKDu case status.

**Table 2.**
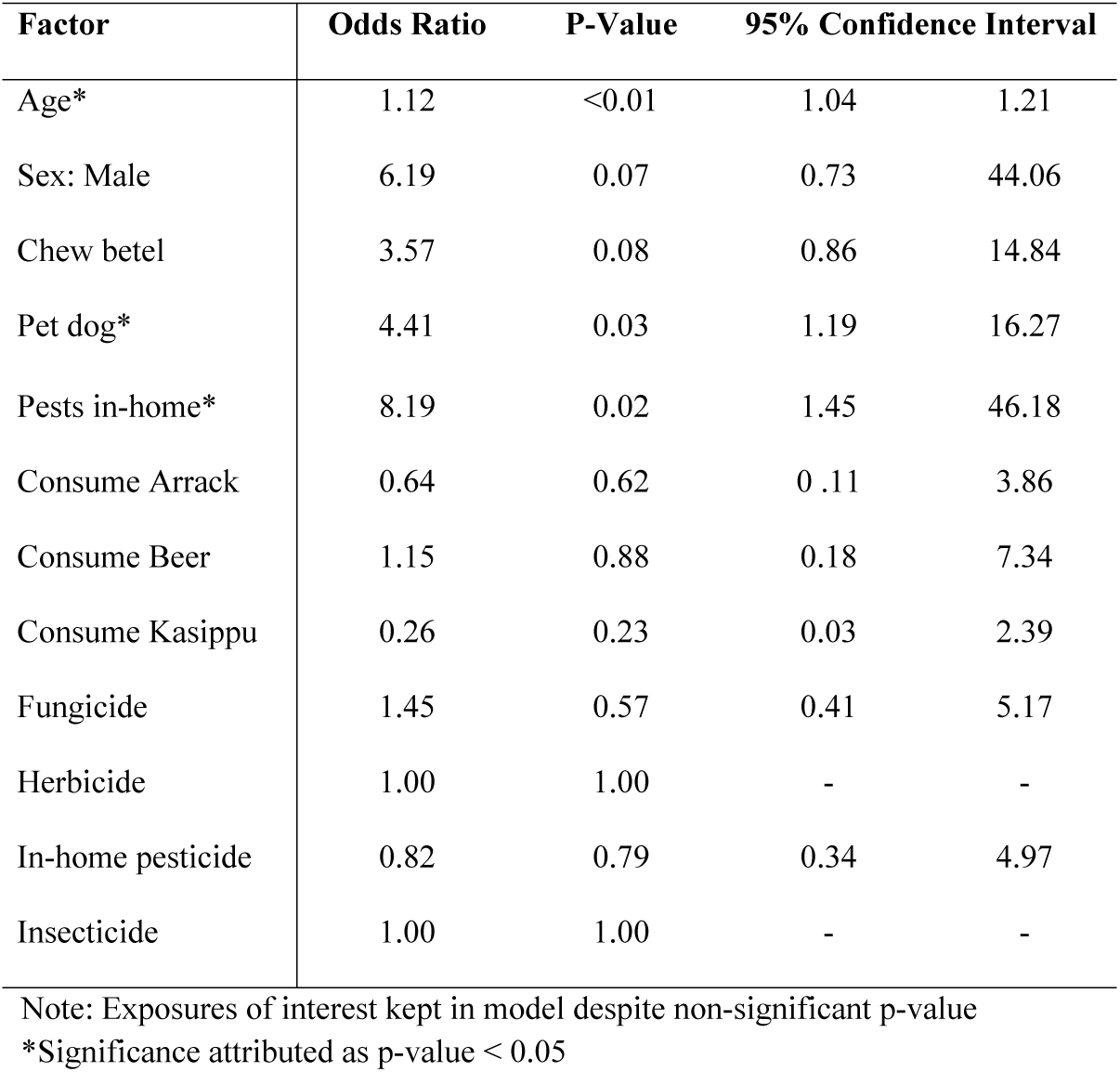
Factors Associated with CKDu including Pesticide and Alcohol Consumption in Sri Lanka.

**Table 3.** Factors Associated with CKDu in Sri Lanka.

| Factor | Odds Ratio | P-Value | 95% Confidence Interval |  |
| --- | --- | --- | --- | --- |
| Chew betel* | 5.95 | 0.002 | 1.878 | 18.856 |
| Pet dog* | 3.515 | 0.012 | 1.312 | 9.42 |
| Treat water* | 3.944 | 0.026 | 1.175 | 13.236 |
| Pests in-home* | 5.708 | 0.009 | 1.538 | 21.182 |
| Age*† | 1.078 | 0.003 | 1.025 | 1.133 |

\*Significance attributed as p-value < 0.05; † Odds ratio per one year age increase

Bivariate analyses for specific types of pesticides and fertilizers used are detailed in Table 4. Before adjustment for age, gender, occupation, and alcohol consumption, usage of a fertilizer (muriate of potash) and an herbicide (glyphosate) were significantly associated with confirmed CKDu cases. However, after adjustment, none of the fertilizers, insecticides, or herbicides reported were found to be significantly associated with the outcome of interest, and so were not included in multivariable models (Table 4).

**Table 4.** Agrochemical Association with CKDu - Crude and Adjusted Odds Ratios

| **Factor** | **Control**<br>**(N=54)**<br>**n(%)** | **Case**<br>**(N=56)**<br>**n(%)** | **Crude Odds**<br>**Ratio (95% CI)** | **Adjusted Odds**<br>**Ratio (95% CI)** |
| --- | --- | --- | --- | --- |
| **Fertilizer** |  |  |  |  |
| Urea | 38 (70%) | 47 (84%) | 2.20 (0.9, 5.5) | 0.92 (0.3, 3.2) |
| Muriate of Potash* | 1 (2%) | 0 (0%) | 3.15 (1.4, 7.4) | 1.86 (0.7, 5.1) |
| Triple Super Phosphate | 11 (20%) | 21 (38%) | 2.35 (1.0, 5.5) | 1.84 (0.7, 5.2) |
| Mud/Manure | 11 (20%) | 5 (9%) | 0.38 (0.1, 1.2) | 0.41 (0.1, 1.5) |
| **Insecticide** |  |  |  |  |
| Carbosulfan | 5 (9%) | 3 (5%) | 0.55 (0.13, 2.44) | 0.48 (0.1, 2.5) |
| Carbofuran | 4 (7%) | 4 (7%) | 0.96 (0.23, 4.06) | 0.47 (0.1, 2.4) |
| Curateer | 7 (13%) | 6 (11%) | 0.81 (0.25, 2.57) | 0.74 (0.2, 2.8) |
| **Herbicide** |  |  |  |  |
| Glyphosate* | 19 (35%) | 32 (57%) | 2.46 (1.14, 5.30) | 1.09 (0.4, 2.8) |
| MCPA | 24 (44%) | 28 (50%) | 1.25 (0.59, 2.65) | 0.92 (0.4, 2.3) |
| DPA | 11 (20%) | 17 (30%) | 1.70 (0.71, 4.08) | 0.88 (0.3, 2.5) |
| Metamifop | 8 (15%) | 8 (14%) | 0.96 (0.33, 2.77) | 1.39 (0.4, 4.9) |

\* $p \leq 0.05$ ; Adjusted by age, gender, farming occupation and alcohol consumption

Figure 2 illustrates the difference between cases and control by type of alcohol reportedly consumed. Cases were more likely to report drinking. The overlap between all three types of alcohol indicates that if one consumes alcohol it is common to drink all three types surveyed. Arrack was the most commonly reported alcohol consumed across cases and controls. Overall reported alcohol consumption was observed among 2.3% of women, while 81.8% of men reported drinking alcohol (two-sided Fisher’s exact: <0.001).

**Figure 2:**
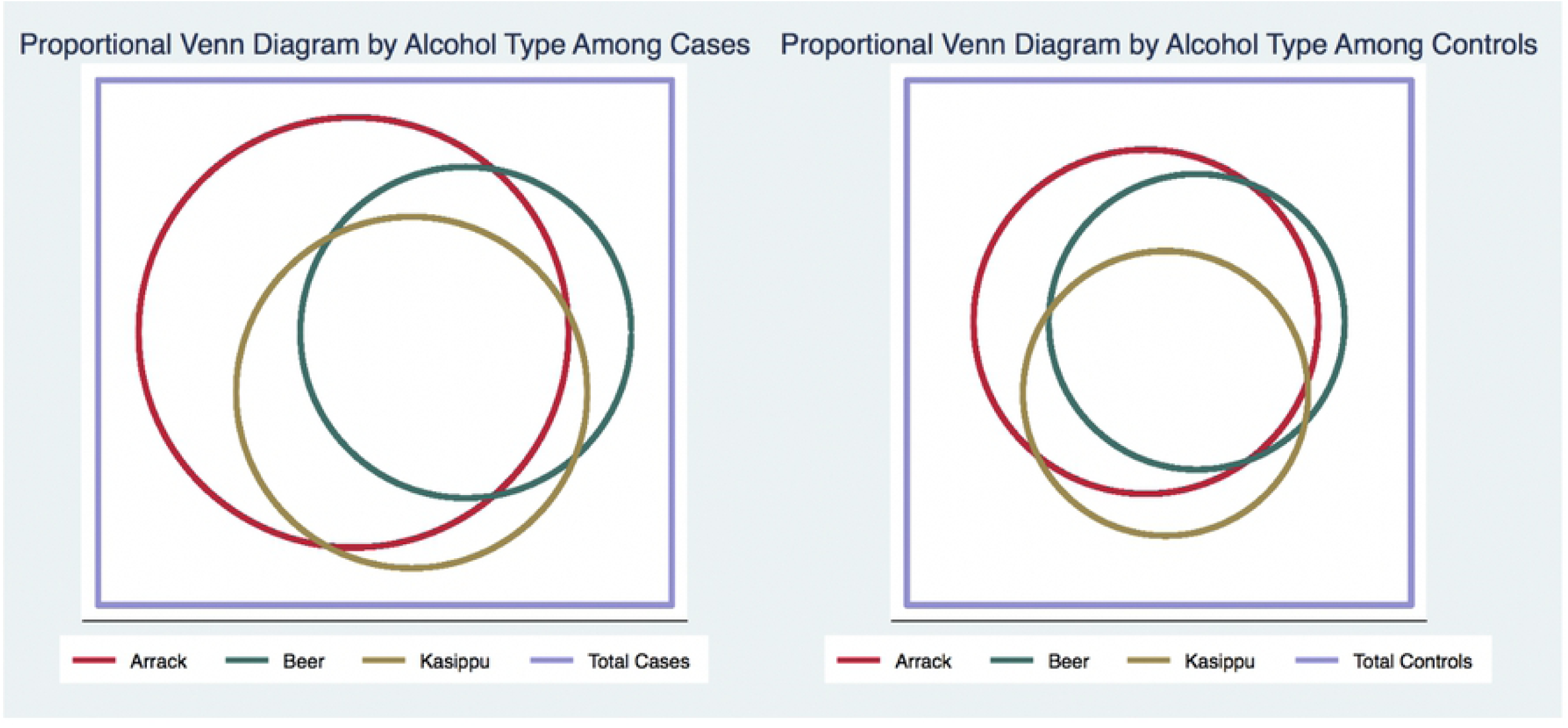
Proportional Venn Diagrams Representing Reported Alcohol Type Consumed by Case-Control Status

## DISCUSSION

There was no significant association detected between CKDu and pesticide exposure nor alcohol consumption. However, there were significant associations identified for chewing betel, owning a pet dog, treatment of drinking water, reporting pests in the home, and age. These significant exposures provide insight into previously unconsidered routes and mechanism for CKDu in addition to potential guidance on how to reduce odds of CKDu diagnosis. Chewing betel could be a risk directly or indirectly through contamination of the traditional chew ingredients or through handling and/or preparation of the betel chew. The association between report of a pet dog could suggest a zoonotic pathway and pests in the home could indicate pest extermination agent risks or a disease reservoir. Treatment of drinking water, especially boiling, may contribute to consumption of aluminum with nephrotoxic consequences. The wide variety of associated exposures suggests that there may be multiple risk factors associated with CKDu, which is consistent with results of previous studies [35, 36].

It is important to note that those cases reporting ever having been diagnosed with diabetes or hypertension were diagnosed after meeting the CKDu case definition. Prevalence of diabetes among controls was high (24.1%) relative to the 2011 and 2030 national estimates of 7.8% and 9.1%, respectively [37]. The questionnaire did not differentiate between type I or type II diabetes mellitus, limiting inferences about reasons for this discrepancy in prevalence. It is possible that changes in meal preparation (bivariate OR = 1.60 95% CI 0.74-3.44) and/or food stuffs available in rural Sri Lanka also play a role in the increased prevalence of diabetes in this control sample. Conversely, diagnosis of hypertension is significantly higher (Fisher’s exact = 0.002) among the cases. However, this is common sequelae of chronic kidney disease [38-40].

Uses of a variety of agrochemicals are common throughout the farming dry zone regions and are often readily available through government subsidies. In our study, only herbicide use was shown to be significant in the bivariate analysis among all insecticide, fungicide, and in-home pesticide parameters. It is possible that exposure to pesticide occurs among those farmers reporting no use of pesticides on their crops through adjacent farm pesticide use in tandem with dynamic environmental factors, i.e. flooding, water source contamination, and winds.

The ingredients used in making kasippu, an illicit locally brewed alcohol, were of special interest in this study, due to prior hypothesis that pesticides are introduced in the brewing process. However, study participants who reported drinking kasippu most often purchased it from other villagers and either did not know the ingredients used or did not want to report drinking kasippu due to it being an illicit form of alcohol in Sri Lanka as well as the perceived cultural stigma for reporting use. Of those that did report drinking kasippu, some reported urea as an ingredient in kasippu production, which could have potentially toxic biologic effects, leading to increased blood urea nitrogen subsequently impacting kidney function [41, 42].

Previous studies have suggested that drinking water quality and contamination may be associated with CKDu [43-45]. Prior studies have identified cases and controls on the basis of groundwater source and found much larger odds of disease in males who drank water from shallow wells, compared to males who drank from natural springs (OR 5.48 95% CI 3.46-8.66) [46, 47]. Similar findings were found for women drinking from shallow wells (OR 4.40 95% CI 2.23-8.68) [46]. Due to the broad application of pesticides in all aspects of farming, potentially nephrotoxic pesticide agents contaminating the drinking water cannot be ruled out. Our findings differ somewhat from these prior studies in the minor difference between 89% (controls) and 93% (cases) reported source of drinking water as a dug well. We however, were not able to compare other sources of drinking water with high confidence and statistical power.

Dug wells are traditional wells often lined with clay brick and may be covered to prevent animals from entering. There were few participants who received water via a tap line, rainwater collectors, or methods other than a dug well. As such, our ability to evaluate drinking water source as a risk factor was limited. In addition, the number of years that participants used different types of drinking water was inconsistently recorded. Treatment of drinking water was found to be a significant risk factor for CKDu. Treatment of water included boiling water (n=41), filtering water (n=59), and traditional methods (n=19). The most common traditional practice for water treatment was the introduction of *Strychnos potatorum* seeds (Sinhalese - ingini seeds) into the water source, as is customary in Sri Lanka and India [48]. One possible risk for developing CKDu related to treating drinking water could be the boiling (n = 41) of water in aluminum vessels [49]. Information regarding the type of cookware used with relation to boiling water was not collected.

Our study found novel risk factors for CKDu in the study region. Results regarding the potential mechanisms of association with CKDu for chewing betel, treatment of drinking water, and having pets or pests are inconclusive. For instance, dogs in Sri Lanka are often community pets and as such, ownership can be difficult to ascertain, although 43% of study participants reported having a pet dog. Additionally, dogs may serve as reservoirs of infectious disease that may lead to renal damage in humans. For example, outbreaks of leptospirosis have been reported during the monsoon season in some CKDu endemic regions [50, 51]. Leptospirosis causes acute kidney failure in dogs, with high mortality rates among those that are untreated or without access to dialysis. Consequently, dogs may potentially shed *Leptospira* spirochetes in their urine. Owners or community individuals may be thus exposed, potentially clearing the infection but suffering renal damage that could later lead to developing CKDu [52]. Current Ministry of Health CKDu diagnosis criteria do not include serology or polymerase chain reaction (PCR) tests for ruling out leptospirosis; patients are asked only to self-report previous history of leptospirosis. As leptospirosis diagnostics continue to improve, future research should consider specimen collection for laboratory confirmation [53-55].

Mammalian pests in households may also be carriers of infectious disease that leads to increased susceptibility to CKDu. There is an emerging hypothesis that Hanta virus could be the possible causative agent for CKDu in Sri Lanka’s dry zone [26, 56]. Humans infected with Hanta virus show clinical signs similar to those of Leptospirosis, and Hanta virus infection in humans was first described in Sri Lanka in the mid-eighties [57]. Rodents act as the reservoir host for Hanta virus and Leptospirosis. Therefore, our study may provide further evidence for the association between infected rodents and prevalence of CKDu in Sri Lanka. Further research should evaluate the incidence of Hanta virus and Leptospirosis in rodents, pets, livestock, and people in CKDu-endemic regions in Sri Lanka.

To identify other disease risks that might be due to close contact with domestic animals or wildlife, data was collected concerning participant observations of illness among domestic animals in the community. There were very few reported, including three observations of ill cattle (2.7%), one goat (0.9%), and one chicken (0.9%) in the month prior to the survey interview. It is possible that the lifespan of dogs and livestock in the community is short due to environmental hazards (disease, trauma, etc.) and animals go missing or die before disease can be detected or transmitted to humans.

Chewing betel was another novel risk factor for CKDu in our study. This practice is quite common among Sri Lankans, and our study found that chewing betel was more common among those reporting farming as an occupation (60%) than other occupations (40%). There is evidence that betel preparations include stimulant properties similar to nicotine, and chewing it routinely can lead to enamel erosion and oral cancer [58-60]. The betel preparation commonly chewed in Sri Lanka is comprised of betel, areca nut, tobacco, and lime. However, betel recipes among farmers in Sri Lanka’s dry zone may contain differing substances compared to preparations in the remainder of the country, which could be associated with CKDu [58, 61, 62]. Individuals that mix and distribute pesticides may be at greater risk, as betel is inserted in the mouth and may be done in the field where hand washing is not possible. In addition, there is evidence that chewing betel increases exposure to arsenic and cadmium, both of which can be nephrotoxic [63].

Additionally, at our sample size (n=110), we do not have the power to detect small differences in effect measure or to generalize to populations outside our study region. Finally, although survey questions were detailed in nature, the question interpretation by the study participant may have varied, leading to inaccurate answers. Due to the condensed nature of the survey timeline, with multiple interviews occurring at one time, investigators could not oversee each individual interview. As such, details pertaining to the number of years when specific pesticides or water sources were used were sometimes incomplete.

This case-control study design was useful for having comparable populations with and without the disease in order to efficiently evaluate past risk factors associated with disease status. In the future, a cohort study would be a useful design to evaluate exposure data, exposure timelines, and incidence rates for CKDu, and we recommend that future studies of CKDu in Sri Lanka be cohort-based, despite the longer follow-up period and greater expense. In addition, a nationwide, coordinated CKDu research consortium spanning all major research institutions would make CKDu research more efficient by standardizing study design and methodologies. This would allow more accurate conclusions to be drawn from studies with clear and consistent case/control definitions and study locations.

While our survey was comprehensive, our study had several limitations, primarily relating to case definition, disease progression, and study design characteristics. Temporal bias might have been introduced since the exact date of CKDu diagnosis was not collected however, the impact of bias on alcohol consumption is believed to be minimal due to a non-significant difference in the mean years since first drink (t = −1.89) and frequency of alcohol consumption (t = −0.49). Furthermore, there was not a significant difference in the number of years farming among those reporting a farming occupation (Fisher’s exact = 0.07) between cases and control.

The appearance and progression of CKDu can involve non-specific symptoms, making the disease challenging to diagnose in the early, pre-clinical stages limiting the population of cases in this study to those that were in advanced stages of the disease. Studies suggest using more sensitive methods for detecting early CKDu, with measurement of microalbuminuria and functional markers such as Cyst C, creatinine or tubular proteins like RBP4, NGAL or KIM would be beneficial in capturing a greater number of early-stage CKDu cases so that exposures can be better assessed and treatment initiated earlier [64-66].

The lack of association between herbicide and CKDu outcome in the adjusted model indicates either that herbicide alone is not responsible for CKDu, or that sufficient detail regarding herbicide use was not captured in the survey. For example, the volume and length of use of pesticides was difficult to assess in an interview setting compared to a household visit, where farmers could reference pesticide receipts or other family members for details they could not recall. More detailed responses regarding usage may have resulted in a difference between cases and controls, which could not be evaluated in this study.

At present, the use of the albumin-creatinine ratio or persistent proteinuria as an initial screening tool is only sensitive at detecting advanced-stage prevalent CKDu cases. This could cause misclassification of the disease outcome if the diagnostic test sensitivity is not high enough to differentiate between early-stage CKDu patients and controls.

## CONCLUSION

In conclusion, this pilot case-control study showed that chewing betel, reporting of in-home pets and pests, treating drinking water, and age were significantly associated with CKDu. These cultural and environmental factors are likely part of a multi-factorial etiology that is challenging to unravel, and that may take years to understand whether preventive measures are effective. Future studies should be cohort in design and focus on further exploring the identified risk factors and their epidemiologic relationships to CKDu, as well as possible interventions to attenuate the incidence of CKDu in Sri Lanka. Potential interventions to be considered based on these findings might include safe home pest control options, testing and treatment for leptospirosis among community dogs, routine chronic kidney disease screening among those in CKDu endemic areas, and education focused on hand hygiene in the field. These findings should be considered as research regarding CKDu in other endemic regions continues.

## DECLARATIONS

### Ethics approval and consent to participate

The study was approved by the University of California, Davis, Institutional Review Board (#762486-2). All participants voluntarily provided written consent prior to questionnaire administration in their preferred language (Sinhala or Tamil).

### Consent for publication

No personal, identifiable information is depicted nor represented.

### Availability of Data and Material

Full questionnaire made available in additional file 1.

### Author’s contributions

All authors listed in this manuscript meet the International Committee of Journal Editors criteria for authorship.

## Acknowledgements

Data collection for this study was made possible through the support provided by the Centre for Research, Education, and Training on Kidney Diseases (CERTkID), namely by the late Mr. Ranjith Mulleriya. Twelve graduate students of University of Peradeniya contributed to the study by interviewing the cases and controls. David Bunn, Ph.D. and Michael Wilkes, M.D. contributed to the researchers’ understanding of the cultural role of alcohol and pesticide use.

## APPENDIX

Table S1: Comprehensive Participant Characteristics by Exposure Category

## SUPPORTING INFORMATION LEGENDS

*Figure 1:* Map of Sri Lanka by district identifying jurisdiction in which survey participant was born. Map produced using QGIS v2.18 (Free Software Foundation Inc. Boston, MA). Jurisdiction boundaries as specified by GADM v2.8, Center for Spatial Statistic, University of California, Davis, California, USA, accessed November 2015. https://gadm.org/data.html.

*Figure 2:* Proportional Venn Diagram produced using Stata IC 14 (Stata Corp., College Station, TX).

Table S1: Characteristics of Population Surveyed: exhaustive list of characteristics among survey participants by exposure category.

